# A time-resolved Förster resonance energy transfer assay to investigate inhibitor binding to ABCG2

**DOI:** 10.1101/2023.10.27.564323

**Authors:** James I. Mitchell-White, Deborah A. Briggs, Sarah J. Mistry, Hannah A Mbiwan, Barrie Kellam, Nicholas D. Holliday, Stephen J. Briddon, Ian D. Kerr

**Author notes:** Corresponding authors: Ian D Kerr; James Mitchell-White.

## Abstract

The human ATP-binding cassette (ABC) transporter, ABCG2 is responsible for multidrug resistance in some tumours. Detailed knowledge of its activity is crucial for understanding drug transport and resistance in cancer, and has implications for wider pharmacokinetics. The binding of substrates and inhibitors is a key stage in the transport cycle of ABCG2. Here, we describe a novel binding assay using a high affinity fluorescent inhibitor based on Ko143 and time-resolved Förster resonance energy transfer (TR-FRET) to measure saturation binding to ABCG2. This binding is displaced by Ko143 and other known ABCG2 ligands, and is sensitive to the addition of AMP-PNP, a non-hydrolysable ATP analogue. This assay complements the arsenal of methods for determining drug:ABCG2 interactions and has the possibility of being adaptable for other multidrug pumps.

**Highlights:**

- ABCG2 is a multidrug pump which moves between states having low or high affinity for substrates and inhibitors
- We introduce a time-resolved Förster resonance energy transfer assay to measure interaction of substrates and inhibitors to ABCG2
- We confirm that NBD dimerization is associated with a switch from a high to a low affinity site for an ABCG2 inhibitor

## Introduction

The distribution and disposition of drugs depends in part on the activity of membrane transporters. The ATP-binding cassette (ABC) superfamily includes several transporters with major roles in the efflux of drugs from cells. The transporters with the broadest specificity are known as the multidrug transporters, which include ABCB1, ABCG2, ABCC1[1, 2], and ABCC5 [3, 4]. ABCG2 was originally discovered through its expression in placenta [5] and in breast tumour cell lines [6] resistant to multiple chemotherapeutic drugs. Beyond its importance in the export of xenobiotics, ABCG2 has a role in urate transport, revealed through the association of ABCG2 polymorphisms with gout [7], and in porphyrin transport[8, 9]. The broad substrate specificity of ABCG2 is revealed by its ability to transport these endogenous compounds but also a range of exogenous molecules including statins, tyrosine kinase inhibitors and other cancer chemotherapy drugs such as mitoxantrone and irinotecan[10]. Understanding and quantifying this polyspecificity remains a key goal in ABCG2 research. Existing assays for ABCG2 activity include determining the effect of transport substrates on ATPase activity (as these are tightly coupled in most ABC transporters) or examining transmembrane transport in either ABCG2-enriched vesicles or live cells. However, none of these assays report quantitatively on the binding event between the transporter and the ligand: to date only radioligand binding assays[11], and microscale thermophoresis[12] have enabled direct determination of ABCG2:ligand interaction but these assays have limitations, such as the need for expensive and hazardous radioactive ligands, or difficulty in understanding the nature of the binding. Given the broad substrate specificity of ABCG2, novel binding assays that can report on transporter pharmacology are a welcome addition to the experimental arsenal.

The potent ABCG2 inhibitor, Ko143 was derived from a fungal toxin, fumitremorgin C[12-14]. It has a high affinity for ABCG2, and binds in a pocket that seems common to other ABCG2 substrates[12, 15-17]. It has also been shown that Ko143 binding tolerates chemical modification[12], making a Ko143 derivative a good candidate for the ligand used in a competitive binding assay.

Förster Resonance Energy Transfer (FRET) based assays have provided rapid and sensitive assays for ligand binding that have been useful for pharmacological studies of receptors[18, 19]. Lanthanide cryptates have long fluorescence lifetimes compared to many other fluorophores, meaning that their signal can be separated by using an integration period longer than the fluorescence lifetimes of other fluorophores. Time resolved FRET (TR-FRET) exploits this to achieve high signal/noise ratios for very sensitive assays. Pharmacological investigation of transporters using TR-FRET is relatively immature, but the advances in receptor studies could serve as a model for transporters.

Here, we demonstrate a sensitive binding assay for ABCG2 based on TR-FRET between a terbium cryptate labelled ABCG2 and a fluorescent derivative of Ko143. Using this assay we determine binding affinity estimates for known ABCG2 substrates. We also demonstrate that this binding assay is sensitive to the conformation of ABCG2 by addition of the non-hydrolysable ATP analogue, AMP-PNP.

## Materials and Methods

### Materials

SNAP-Lumi4-Tb, and LabMed buffer were purchased from CisBio. All other materials purchased from Sigma-Aldrich (UK). HEK293S cells were provided by Prof. Dmitry Veprintsev, University of Nottingham. Chemicals and solvents of analytical and HPLC grade were purchased from commercial suppliers and used without further purification. BODIPY-630/650-X-SE, BODIPY-FL-X- SE and BODIPY-FL-SE were purchased from Molecular Probes (Thermo Fisher Scientific).

### Methods

#### Synthesis and chemical analysis of Ko143-X-BY630

All reactions were carried out at ambient temperature unless otherwise stated, with full details provided in the Supplementary Methods. Reactions were monitored by thin-layer chromatography on commercially available silica pre-coated aluminium-backed plates (Merck Kieselgel 60 F254). Visualisation was under UV light (254 nm and 366 nm). PTLC purification was carried out using commercially available pre-coated glass backed silica plates (Analtech, 20 x 20 cm, silica gel 60 matrix). NMR spectra were recorded on a Bruker-AV 400. ^1^H spectra were recorded at 400.13 Hz and ^13^C NMR spectra at 101.62 Hz. All ^13^C NMR are ^1^H broadband decoupled. Solvents used for NMR analysis (reference peaks listed) were CDCl_3_ supplied by Cambridge Isotope Laboratories Inc., (δ_H_ = 7.26 ppm, δ_C_ = 77.16) and CD_3_OD supplied by VWR (δ_H_ = 3.31 ppm and δ_C_ = 49.00). Chemical shifts (δ) were recorded in parts per million (ppm) and coupling constants are recorded in Hz. Processing of the NMR data was carried out using the NMR software Topspin 3.0. LC-MS spectra were recorded on a Shimadzu UFLCXR system coupled to an Applied Biosystems API2000 and visualised at 254 nm (channel 1) and 220 nm (channel 2). LCMS was carried out using a Phenomenex Gemini-NX C18 110A, column (50 mm × 2 mm x 3 μm) at a flow rate 0.5 mL/min over a 5 min period. All high-resolution mass spectra (HRMS) were recorded on a Bruker microTOF mass spectrometer using MS electrospray ionization operating in positive ion mode. RP-HPLC was performed on a Waters 515 LC system and monitored using a Waters 996 photodiode array detector at wavelengths between 190 and 800 nm. Spectra were analysed using Millenium 32 software. Semi-preparative HPLC was performed using YMC-Pack C8 column (150 mm × 10 mm × 5 μm) at a flow rate of 5.0 mL/min using a gradient method of 50-60% B over 12 minutes (Solvent A = 0.01% formic acid in H_2_O, solvent B = 0.01% formic acid in CH_3_CN. Analytical RP-HPLC was performed using a YMC-Pack C8 column (150 mm × 4.6 mm × 5 μm) at a flow rate of 1.0 mL/min. The retention time of the final product is reported using a gradient method of 5-95% solvent B in solvent A over 25 minutes. (Solvent A = 0.01% formic acid in H_2_O, solvent B = 0.01% formic acid in CH_3_CN).

#### Lumi4-Tb labelled ABCG2

A human ABCG2 (NCBI Gene 9429) construct was engineered with sequential N-terminal TwinStrep tag and SNAP-tag, and this was cloned into the pcDNA3.1/Zeo(+) plasmid. HEK293S cells were transfected with this plasmid as in previous ABCG2 studies[20] and cells expressing TwinStrep-SNAP-ABCG2 (293S-SNAP-G2) were selected and maintained using zeocin at concentrations of 40 and 200 μg mL^-1^ respectively. Protein expression was confirmed by western blot using the BXP-21 antibody, and Ko143-inhibitable efflux of mitoxantrone was confirmed by flow cytometry as described previously[21, 22]. Briefly, cells are treated with 10 μM mitoxantrone with or without 1 μM Ko143, then with only Ko143 or a solvent (DMSO) control. Following centrifugation and washing, cells were analysed in a MoFlo Astrios flow cytometer. The m itoxantrone fluorescence of at least 10,000 cells was determined. ABCG2 function was measured as a rightward shift in the 645 nm channel.

For TwinStrep-SNAP-ABCG2 labelling, 293S-SNAP-G2 cells were seeded at 2.5 x 10^5^ cells mL^-1^ in T-75 flasks in 40 mL DMEM + 10% FBS, supplemented with 0.1% w/v pluronic f-68. Flasks were stood vertically on a shaker at 200 revolutions per minute (RPM) until their density exceeded 2.5 x 10^6^ cells mL^-1^. 24 hours before harvesting 10 mM sodium butyrate was added to enhance protein expression[23-25]. Cells were harvested by centrifugation at 300 g for 15 minutes and lysed by nitrogen cavitation for 20 minutes at 1000 psi before centrifugation at 300 g, 4 °C to collect debris, followed by centrifugation at 100,000 g, 4 °C to collect a membrane pellet. Membrane pellets were resuspended by serial passage through 21g needles in 10 mM Tris-HCl, pH8 and 250 mM sucrose for storage at -80 °C.

Membrane pellets were resuspended at 100 mg mL^-1^ wet weight of membranes in LabMed buffer containing 100 nM SNAP-Lumi4-Tb, and incubated at room temperature for 1 hour, rotating on a daisy wheel. Membranes were pelleted by centrifugation at 20,000 g for 20 minutes, 4 °C then washed in 50 mM Tris-HCl, pH 8 before resuspending in 50 mM Tris-HCl, pH 8 at 1 mg mL^-1^ wet weight of membranes. Aliquots of labelled membranes could be stored at -80 °C for at least 6 months without loss of functionality.

#### ABCG2 inhibition assay

HEK293S cells, and 293S-SNAP-G2 were seeded in a clear-bottom, black well 96-well plate at 10,000 cells/well 24 hours before assay. Media was exchanged for Hank’s Balanced Salt Solution (HBSS) containing 3 μM Hoechst 33342, with or without 1 μM Ko143 or Ko143-X-BY630. The cells were incubated for 45 minutes, then exchanged for HBSS containing the previously administered inhibitor or a solvent control for 45 minutes. Incubations were carried out at 37 °C. All wells were washed with HBSS and the fluorescence at 350/460nm and 620/660nm (excitation/emission) was measured using a BMG CLARIOstar to measure H33342 and Ko143-X-BY630 fluorescence respectively. Data for each repeat were normalized by subtracting the fluorescence of control cells, then dividing by the mean fluorescence of HEK293S cells with added Hoechst 33342 or Ko143-X- BY630 respectively.

#### Saturation binding TR-FRET assay

Ko143-X-BY630 was prepared from DMSO stock solution at a range of concentrations in LabMed buffer. A Ko143 solution yielding an assay concentration of 30 μM was prepared in LabMed with pluronic F-127 to yield a final assay concentration of 0.02% (w/v). This was mixed with 20 μg membranes as prepared above to a final assay volume of 40 μL. This was allowed to equilibrate for 30 minutes at room temperature before measurement of fluorescence with an excitation of 337 nm, recording emission at 620 nm for donor (Tb) fluorescence, and 665 nm for FRET on a BMG PHERAstar. Each reading was ten excitation flashes, recording an integration period of 60 to 400 μs for emission. Parallel solvent-control assays were performed at the maximum concentration of DMSO achieved.

#### Competitive binding TR-FRET assay

For competition binding assays, ABCG2 ligands were prepared over concentration ranges according to their anticipated affinities. These were added to membranes simultaneously with Ko143-X-BY630 (50 nM). Following incubation at room temperature for 30 minutes, fluorescence was measured as for the saturation binding assay above. A sample with 30 μM Ko143 was also measured to estimate values for 100% inhibition.

#### Analysis

Unless otherwise specified, reported results are the mean ± standard deviation of at least three independent experiments each with at least technical triplicates. FRET ratio and specific binding were calculated using GraphPad Prism 9.

For TR-FRET experiments, FRET ratio was calculated as

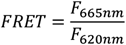

In saturation binding experiments, FRET ratios were fitted to a one site binding model, fitting both total and non-specific binding to a model with a unit Hill slope:

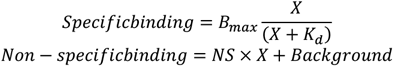

The concentration curve with added Ko143 was fit to the non-specific binding model, and the concentration curve without added Ko143 was fit as the sum of specific and non-specific binding.

In competition binding experiments, TR-FRET ratios were calculated, then these values were normalized to percentage of binding inhibition with 0% as the competing ligand-free reading, and 30 μM Ko143 as 100% inhibition. For each ligand, an IC50 was estimated by fitting these values to a model with unit Hill slope:

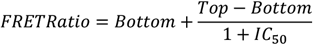

The corresponding *K*_*i*_was calculated using the Cheng-Prusoff equation[26]:

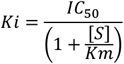

## Results

### SNAP-tagged ABCG2

Initially a tagged ABCG2 protein was characterized for functionality. A plasmid encoding a TwinStrep-SNAP-tagged ABCG2 was constructed and expressed robustly following zeocin selection in HEK293S cells (Figure 1a). This construct was functional at the plasma membrane and transported mitoxantrone comparably to another fully-functional ABCG2 construct, with a hexahistidine tag, as can be seen in Figure 1b. This afforded confidence that the tagging of ABCG2 at its N-terminus with a Twin-Strep and SNAP tag does not impair either localization or function, in agreement with other cell-based functional studies of ABCG2[27].

**Figure 1:**
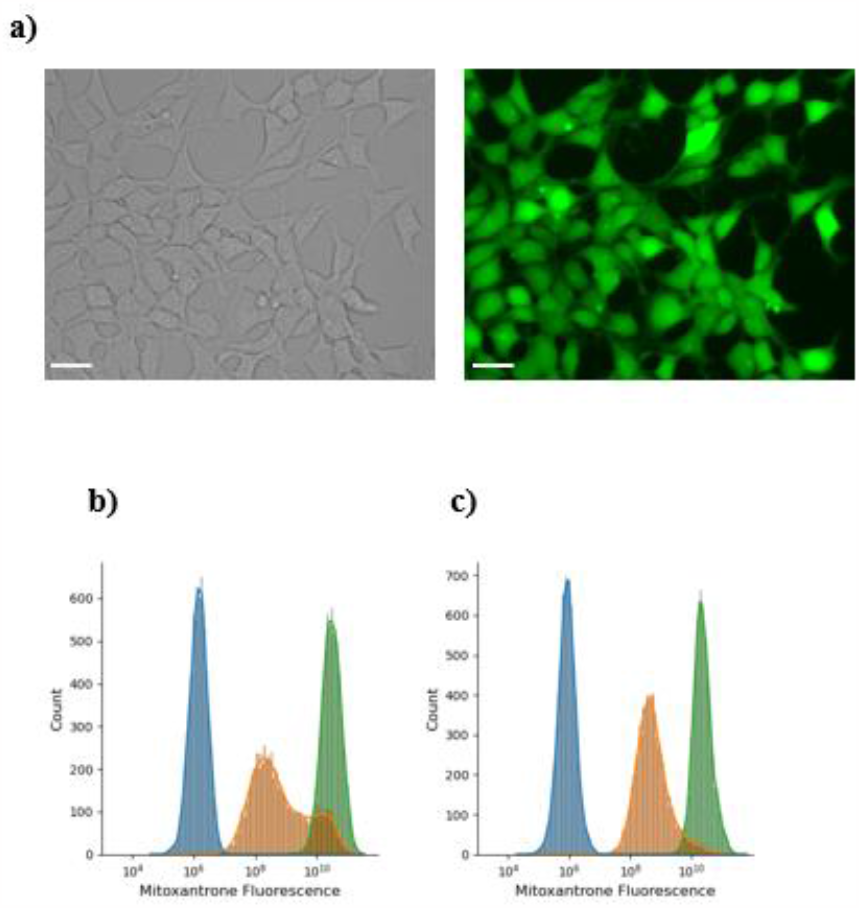
Expression and function of TwinStrep-SNAP-ABCGZ. **a)** Expression of SNAP-ABCG2 in HEK293S cells detected by SNAP-Cell Oregon Green labelling. Scale bar 10 μm b) Transport of mitoxantrone by SNAP-ABCG2. Efflux of mitoxantrone by ABCG2 was measured by flow cytometry. Mitoxantrone fluorescence in a vehicle control (blue), a mitoxantrone treated sample (orange) and with inhibition by Ko143 (green) were compared in **b)** HEK293T expressing His-ABCG2 and **c)** HEK293S expressing SNAP-ABCG2. Data are representative of at least 3 independent repeats.

### A fluorescent Ko143 derivative

The fumitremorgin C analogue Ko143 is a high affinity and selective ABCG2 inhibitor, which displays more than 200-fold selectivity over ABCB1 and ABCC1 [13]. This ligand was used as the starting point for the development of a fluorescent ligand for ABCG2 for use in a TR-FRET assay. Fluorescent ligands typically consist of a target binding moiety, a linker and a fluorophore. To connect the target binding moiety (Ko143) to the fluorophore (BODIPY-630/650-X) a short ethylene diamine linker was employed. The red-emitting BODIPY-630/650-X fluorophore is a frequently utilised fluorescent dye for fluorescence binding assays, offering good optical properties including a high quantum yield and a low extinction coefficient.

To synthesise the fluorescent ligand, the tert-butyl ester of commercially available Ko143 (compound **1**, Figure 2a) was firstly converted to the corresponding carboxylic acid **2** in a 66% yield after aqueous work-up. The following step was an EDC.HCl coupling of N-Boc ethylene diamine to the carboxylic acid which gave compound **3** in a 55% yield after purification by PTLC and removal of excess ethylene diamine using a PS-benzaldehyde resin. Boc deprotection of the amine was carried out in 2M HCl in 1,4-dioxane which gave compound **4** in a quantitative yield. The amine hydrochloride salt was used directly in the BODIPY-630/650-X coupling step without purification. The final fluorescent ligand **5** was obtained in a 9% yield following purification by PTLC and semi-preparative HPLC.

**Figure 2:**
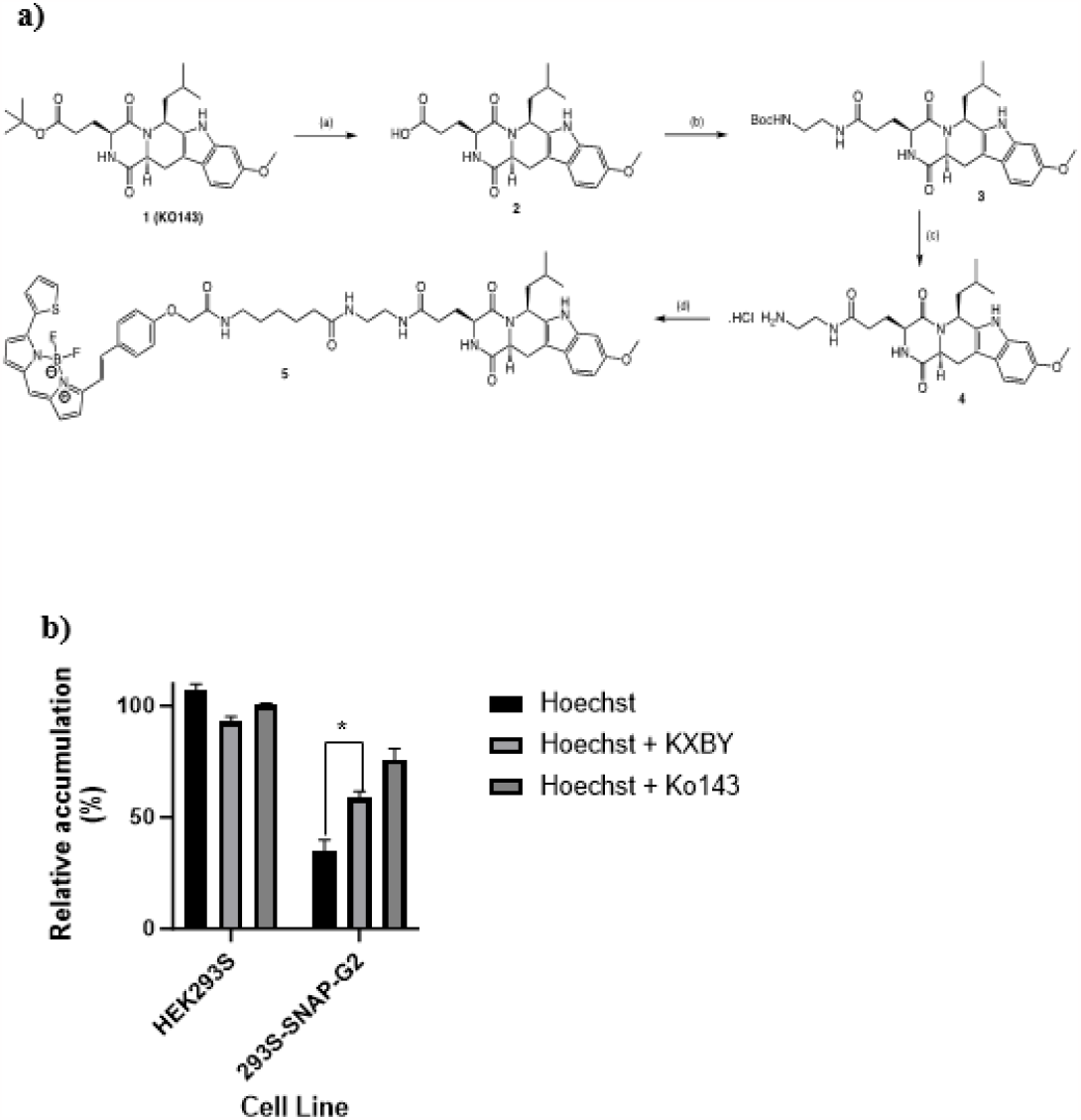
A fluorescent Ko143 retains potential to inhibit transport by ABCG2. **a)** Synthesis of Ko143-X-BY63O. **Reagents & Conditions:** (step a) IFA,DCXL rt, 3 hrs. 5.8 at. 66%. (b) Boc ethylene diamine, EDC.HC1. HOBt, DIPEA. DXIF, rt. 6 days 43 mg, 55%. (c) 4 M HC1 in dioxane, rt, 3 hrs. quantitative. (d) BODIPY-X-630/650-SE. DIPEA, DCM, DMF, rt, 24 hours, 0.2 mg; final yield 8.7 % **b)** Transport of Hoechst 33342 is inhibited by Ko143-X-BY630. HEK293S cells or 293S-SNAP-ABCG2 were incubated with H33342 in the presence or absence of unmodified Ko143 or Ko143-X-BY630. Cellular accumulation ofH33342 was determined by a fluorescence plate reader and is expressed as a percentage of staining of HEK293S cells. Values represent the mean and standard error of three independent experiments. Ko143-X-BY630 causes an increase in H33342 fluorescence (Difference between Control and +Ko143-X-BY630 significant, p = 0 016).

### Functional Characterization

To determine whether Ko143-X-BY630 retained the ability to inhibit transport by ABCG2, whole-cell transport assays of Hoechst 33342 were conducted in untransfected HEK293S cells and those expressing Twin-Strep-SNAP-ABCG2. Hoechst 33342 was selected as a test substrate as its fluorescence excitation and emission does not overlap significantly with those of the BODIPY 630/650 fluorophore. As shown in figure 2b, addition of 1 μM of Ko143-X-BY630 to cells expressing ABCG2 resulted in a significant Hoechst 33342 accumulation, but had no effect on untransfected cells, in a manner similar to the unmodified inhibitor Ko143. We also ruled out that derivatizing Ko143 with BY-630 altered it from an inhibitor to a transport substrate by determining its own accumulation in untransfected and ABCG2-expressing cells with and without the addition of Ko143. The expression of ABCG2 or its inhibition by Ko143 made no difference to the cellular fluorescence intensity of Ko143-X-BY630 (supplementary figure S2) confirming that Ko143-X- BY630 is not a transport substrate for ABCG2.

### Ko143-X-BY630 binds to ABCG2 with high affinity

Once Ko143-X-BY630’s inhibitory activity in cells had been confirmed, its binding affinity to ABCG2 was established by saturation binding using TR-FRET. Ko143-X-BY630 was added to membranes from 293S-SNAP-G2 labelled with Lumi4^®^-Tb terbium cryptate. In order to separate ABCG2 specific binding from non-specific binding, Ko143 was added to the same concentration series, and the FRET signal subtracted. As shown in figure 3, the specific binding follows a one-site binding model. The pK_D_ for Ko143-X-BY630 was 7.79 ± 0.04 (mean ± SEM, mean K_D_ 16.5 nM).

**Figure 3:**
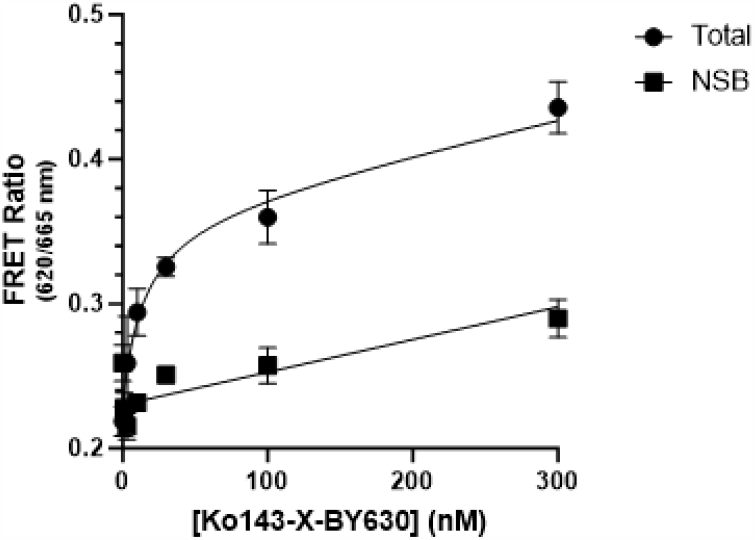
Saturation binding of Ko143-X-BY6J0 to ABCG2. Membrane suspensions with Tb-labelled ABCG2 were incubated with Ko143-X-BY630, with and without unlabelled Ko143 to isolate specific binding, and TR-FRET was recorded for a range of Ko143-X-BY630 concentrations between 1 and 300 nM. A representative concentration-response carte is shown as the mean 1 SEM of three replicates, fitting both total and non-specific binding, assuming a single binding site. Analysis of 3 independent repeats allowed for a pKD for Ko143-X-BY630 of 7.79 ± 0.04 to be determined. K_D_ 16.5 nM.

### Ko143 can displace Ko143-X-BY630 from ABCG2

To demonstrate displacement of Ko143-X-BY630 by Ko143, a range of Ko143 concentrations were added to Tb-ABCG2 membranes and 50 nM Ko143-X-BY630 (figure 4a). An average K_I_ of 49.9 nM was estimated from these experiments, suggests that the binding of the fluorescent derivative is little affected by the addition of the BODIPY moiety. Similarly, this K_I_ compares well with literature values for the IC_50_ of Ko143 in the as a transport inhibitor (128 nM [28]) and with a K_D_ for a fluorescent Ko143 derivative determined by microscale thermophoresis (4.4 nM[12]).

**Figure 4:**
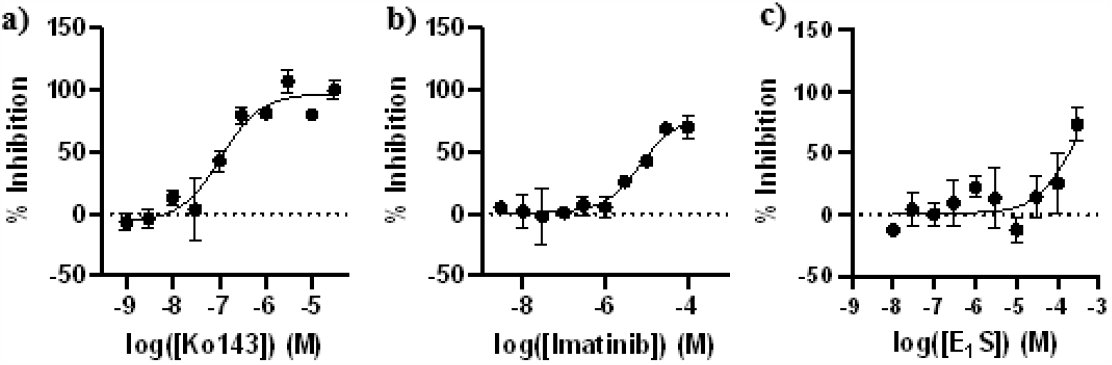
Displacement of Kb143-X-BYtf30 by ABCG2 ligands. Membrane suspensions with Tb-labelled ABCG2 were incubated with 50 nM Ko143-X-BY630 and a range of concentrations of each Iigand to measure the reduction in FRET signal, as a measure of displacement ofKp143-X-BY630. A representative concentration-response curve is shown as the mean ± SEM of three replicates. Analysis of multiple independent repents allowed calculations of IC_50_ and K _1_ values of. **a) Ko143** IC_50_ = 200 ± 74.7 nM, K _1_ = 49.9 nM, pK _i_ = 7.38 ± 0.26. **b) Imatinib** IC_50_ = 5.17 ± 1.95 μM, K _i_ = 1.28 μM, pK _i_ = 6.15 α 0.60. **c) Estrone 3-sulfate** IC _50_ = 336 ± 141 μM, K _i_ = 79.1 μM. pK _i_ = 4.15 ± 0.20. K _i_ obtained from IC _50_ and Cheng-Prusoff correction.

### Competitive ABCG2 binding can detect ABCG2 ligands

Having characterised the binding of Ko143-X-BY630 to ABCG2, its use in a competitive binding assay to characterise known ABCG2 substrates was demonstrated. Estrone 3-sulfate (E_1_S) is a canonical ABCG2 substrate, and has been resolved in a cryo-EM structure of ABCG2[16, 29]. Another ligand that has been used in structural studies is the tyrosine kinase inhibitor, imatinib, which acts as an ABCG2 inhibitor[17, 30]. Both substrates are shown to bind in the same cavity as Ko143. The competition of these ligands with Ko143-X-BY630 was used to find their affinity for ABCG2, shown in figure 4b and 4c. The K_i_ for imatinib was determined as 1.28 μM, and a maximum inhibition of binding of 93%. Estrone 3-sulphate inhibited to a similar extent at the highest concentration. Based on maximal inhibition of 100%, the K_I_ of estrone 3-sulphate was estimated as 79.1 μM.

### Affinity of ABCG2 for Ko143-X-BY630 is reduced in an ATP-bound state

Structural studies have shown that ABCG2 adopts an inward-facing conformation in the absence of ATP[17], and upon ATP binding, ABCG2 adopts an outward-facing conformation[16]. It has also been demonstrated that resetting ABCG2 to an inward-facing conformation is likely to be the step of the cycle dependent on ATP hydrolysis[31, 32]. In many ABC transporters this outward-facing state can be achieved by mutation of the catalytic glutamate to glutamine (E211Q in ABCG2) or by the addition of saturating concentrations of a non-hydrolysable ATP analogue AMP-PNP. This TR- FRET binding assay allows the affinity of different states of ABCG2 to be determined. Adding AMP-PNP increased the Kd approximately 10-fold to 114 nM (figure 5, difference of pKd significant by t-test, p = 0.004).

**Figure 5:**
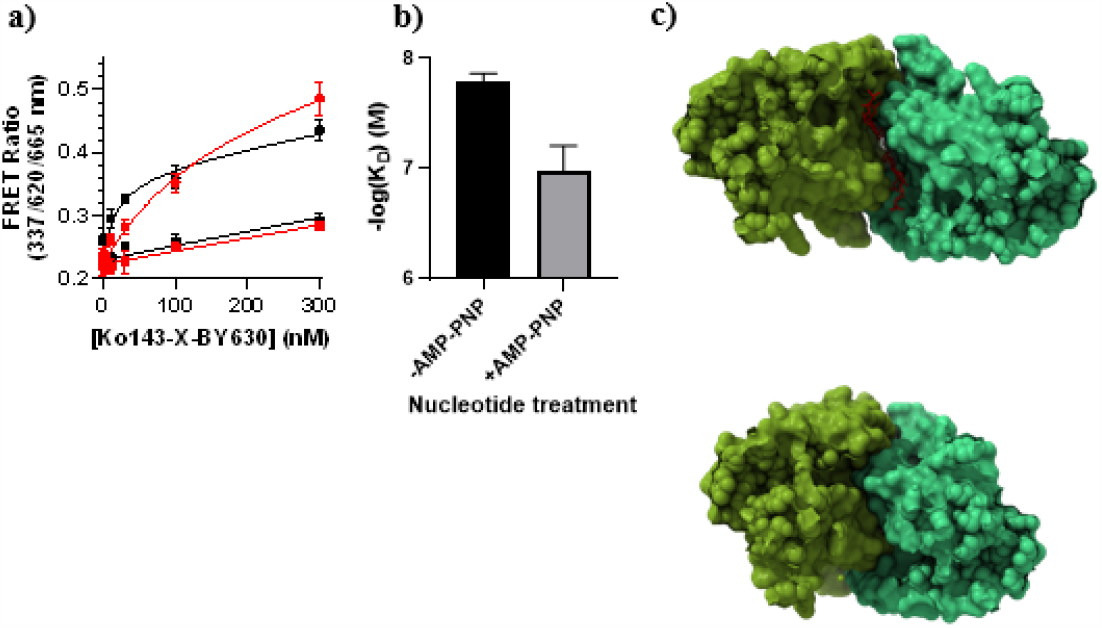
AMP-PNP decreases affinity of Ko143-X-BY630 for ABCG2. **a)** Effect of AMP-PNP upon concentration-response curve of Ko143-X-BY630. Binding of Ko143-X-BY630 to Tb-labelled AJ3CG2 was measured by TR-FRET in the absence (black), and presence (red) of 20 mM AMP-PNP. Total binding represented by circles, non-specific binding by squares. Representative curves shown. **b)** Comparison of treatments. From mean of three independent experiments. K _D_ with AMP-PNP = 114.09 α 42 nM. Difference between pK_D_, significant (p = 0.004) **c)** Binding pocket changes upon AMP-PNP addition. Structures of ABCG2 without bound nucleotide (6ETI upper) and with bound nucleotide (6HBU lower) bound nucleotide were aligned. Structures are shown Horn the cytosolic face, with the NBDs emitted, and Ko143 derivative MZ29 shown in red.

## Discussion

Transporter function is commonly assessed by the determination of transmembrane substrate flux. These assays, frequently reliant on fluorescent substrates, provide data on the overall function of the transporter, rather than individual steps in the transport pathway. The use of fluorescent ligands offers many advantages over the use of e.g. radioligands, such as safety, affordability, and applications to a broader range of techniques[33, 34].

Transport is the sum of a several component events including substrate binding, substrate occlusion and substrate release[35]. For a multidrug efflux pump, the initial events are a membrane partitioning event (as substrates are principally hydrophobic) and a protein interaction event. For ABCG2 this initial drug binding occurs at a higher affinity inward-facing conformation in which the NBDs are ATP-free[32, 36, 37]. Following substrate and ATP binding there is a conformational change to a lower affinity, outward-facing conformation. Binding assays are required to help us understand this initial interaction and the assay described here enables us to analyse ligand binding to the high affinity inward facing state of ABCG2.

In this work, we demonstrated a rapid, miniaturized TR-FRET assay for the binding of substrates to cavity 1 of ABCG2. The assay has potential for high-throughput screening and direct evaluation of ligand affinities and works with ABCG2 in a membrane fraction, thus complementing transport assays in whole cells[31] and binding assays to purified protein[20, 38]. The assay uses a novel, fluorescent analogue of the ABCG2 inhibitor, Ko143. Ko143 was chosen as a basis upon which to build a ligand due firstly to its high affinity, which enables a low assay concentration, minimising the effects of non-specific binding in the assay, and secondly because there is evidence in the literature that Ko143 can be derivatised to be fluorescent without loss of binding to and inhibition of ABCG2[12].

Subsequent structures with bound ABCG2 transport substrates (e.g. mitoxantrone, imatinib, and estrone 3-sulphate (E_1_S)) have confirmed these all bind at cavity 1[15-17, 29]. Additionally, specific interactions with residues in the binding cavity have been shown to be vital for transport, particularly stacking interactions with F439[14].Given these structural data, it seems likely that binding to cavity 1 is a step required for transport by ABCG2. Pharmacological studies demonstrating multiple binding sites in ABCG2[11, 21] may be reconciled with these structural data if they are detecting interaction of drugs at other surface-localized “access” sites as has been observed for other multidrug pumps such as ABCB1[39].

The assay demonstrated here also enables analysis of different conformations of ABCG2. It is sensitive to the presence of the non-hydrolysable ATP analogue, AMP-PNP, showing that a nucleotide-bound ABCG2 adopts a lower affinity conformation. Structurally, the outward facing conformation of ABCG2 has a much narrower binding cavity resulting in the loss of specific interactions known to be vital for transport (figure 5c comparison of cavity 1 in the nucleotide free and bound structures of ABCG2). Our demonstration of a 10-fold reduction in affinity for Ko143-X- BY630 in the AMP-PNP bound state provides more evidence for the hypothesis that ATP binding is responsible for affinity switches in ABCG2[31, 32] and thus that ATP hydrolysis is responsible for transporter resetting to the inward facing conformation.

Understanding the characteristics of multidrug transporters is vital for understanding multidrug resistance. Having tools to explore the binding of substrates to transporters would improve the understanding of their structure-activity relationships, and the broader role of transporters in pharmacokinetics and metabolism. The present assay contributes to the tools available for studying ABCG2 and could be broadened to other MDR pumps [20, 31, 38].

## Abbreviations

ABC: – ATP binding cassette;
FRET: –Forster resonance energy transfer;
TR: – time- resolved;
HEK: – human embryonic kidney;
DMSO: – dimethylsulfoxide;
DMEM: – Dulbecco’s Modified Eagle’s Medium;
FBS: – fetal bovine serum;
HBSS: - Hank’s Balanced Salt Solution;
H33342: – Hoechst 33342;
BODIPY-630/650-X-SE,: 6-(((4,4-difluoro-5-(2-thienyl)-4-bora-3a,4a-diaza-s-indacene-3-yl)- styryloxy)acetyl)aminohexanoic acid succinimidyl ester
BODIPY-FL-X-SE,: 6-((4,4-difluoro-5,7-dimethyl-4- bora-3a,4a-diaza-s-indacene-3-propionyl)amino)hexanoic acid succinimidyl ester
BODIPY-FL-SE,: [1-[3-[5- [(3,5-dimethyl-2*H*-pyrrol-2-ylidene-κ-*N*)methyl]-1*H*-pyrrol-2-yl-κ-*N*]-1-oxopropoxy]-2,5- pyrrolidinedionato]difluoro-,(T-4)-Boron; DCM, dichloromethane
DIPEA,: *N,N*-diisopropylethylenediamine
DMF,: *N,N*-dimethylformamide
EDC.HCl,: *N*-(3-Dimethylaminopropyl)-*N′*-ethylcarbodiimide hydrochloride
HOBt,: 1-Hydroxybenzotriazole hydrate
PTLC,: Preparative Thin Layer Chromatography;
TFA,: trifluoroacetic acid

## Author Contributions

Conceptualization: JIMW, IDK, NDH, SJB

Investigation: JIMW, DAB, IDK, HAM, SJM, BK

Formal analysis and visualization: JIMW

Writing – original draft: JIMW, IDK

Writing – review and editing: JIMW, IDK, NDH, SJB, SJM, BK

## Acknowledgements

We thank Ella Hutchison for technical assistance with some preliminary experiments.

## Funding

This work was supported by Biotechnology and Biological Sciences Research Council [grant number: BB/S001611/1] to IDK, SJB and NDH. HAM was supported by BBSRC Doctoral Training Grant [BB/M008770/1]

## Supplementary Methods

### Full chemical synthesis of Ko143-X-BY630

The following abbreviations are used to described signal shapes and multiplicities; singlet (s), doublet (d), triplet (t), quadruplet (q), broad (br), dd (doublet of doublets), ddd (double doublet of doublets), dtd (double triplet of doublets) and multiplet (m).

**(3*S***,**6*S***,**12a*S*)-1**,**2**,**3**,**4**,**6**,**7**,**12**,**12a-Octahydro-9-methoxy-6-(2-methylpropyl)-1**,**4- dioxopyrazino[1’**,**2’:1**,**6]pyrido[3**,**4-b]indole-3-propanoic acid (compound 2; figure 2a)**

To *tert-*Butyl (3*S*,6*S*,12a*S*)-1,2,3,4,6,7,12,12a-Octahydro-9-methoxy-6-(2-methylpropyl)-1,4- dioxopyrazino[1’,2’:1,6]pyrido[3,4-*b*]indole-3-propanoate (Ko143) (10.0 mg, 21.3 μmol) in DCM (2.0 mL), was added TFA (450 μL). The mixture was stirred at room temperature for 3 hours and the solvent removed under vacuum. The residue was redissolved in DCM (5 mL) and the product extracted with sat. Na_2_CO_3(aq)_, (3 x 5 mL). The aqueous phase was adjusted to pH 4 with acetic acid and then extracted with DCM (3 x 5 mL). The combined organic phases were combined and dried over MgSO_4_ and the solvent removed under vacuum to afford the desired product as a white crystalline solid (5.8 mg, 66%). LCMS *m/z* calc. for C_22_H_28_N_3_O_5_ [M+H]^+^ 414.2; found; 414.2, *t*_R_ = 2.62 min.

***tert*-Butyl (2-(3-((3*S***,**6*S***,**12a*S*)-6-isobutyl-9-methoxy-1**,**4-dioxo-1**,**2**,**3**,**4**,**6**,**7**,**12**,**12a- octahydropyrazino[1’**,**2’:1**,**6]pyrido[3**,**4-*b*]indol-3-yl)propanamido)ethyl)carbamate (compound 3, figure 2a)**

To a solution of compound **2** (5.8 mg, 14.0 ſmol) in DMF (1.0 mL), EDC.HCl (3.2 mg, 16.8 ſmol)), HOBt (8.9 mg, 65.9 ſmol)) and DIPEA (12.2 ſL, 70.0 ſmol)) were added. The mixture was stirred for 10 minutes and *N*- Boc ethylene diamine (6.7 ſL, 42.1 ſmol), was added. The mixture was stirred at room temperature for 6 days with further additions of DIPEA and EDC.HCl (after 24 hours (1 eq), 5 days (0.5 eq) & 6 days (0.2 eq)). The solvent was removed, and the reaction mixture was purified by PTLC (5% MeOH in DCM). Remaining Boc- ethylene diamine was removed with PS-Benzaldehyde resin to afford the title product as a white solid (4.3 mg, 55%). LCMS *m/z* calc for C_29_H_42_N_5_O_6_ [M+H]+ 556.3; found 556.4, t_R_ = 2.87. ^1^H NMR (400 MHz, CDCl_3_) δ 8.08 (s, 1H), 7.71 (s, 1H), 7.43 (d, *J* = 8.6 Hz, 1H), 6.89 (d, *J* = 2.2 Hz, 1H), 6.82 (dd, *J* = 8.6, 2.2 Hz, 1H), 6.59 (s, 1H), 5.45 (dd, *J* = 9.3, 4.2 Hz, 1H), 5.01 (t, *J* = 6.0 Hz, 1H), 4.03 (ddd, *J* = 10.3, 7.7, 5.3 Hz, 3H), 3.85 (s, 3H), 3.51 (dd, *J* = 15.8, 4.9 Hz, 1H), 3.38 (q, *J* = 5.5 Hz, 2H), 3.28 (d, *J* = 5.9 Hz, 2H), 3.04 (dd, *J* = 15.8, 11.7 Hz, 1H), 2.54 – 2.36 (m, 3H), 2.29 (dd, *J* = 9.4, 6.2 Hz, 1H), 1.78 – 1.49 (m, 4H), 1.44 (s, 9H), 1.04 (d, *J* = 6.5 Hz, 3H), 0.82 (d, *J* = 6.3 Hz, 3H). ^13^C NMR (101 MHz, CDCl_3_) δ 173.7, 169.9, 168.8, 161.3, 156.6, 136.7, 133.2, 120.8, 118.9, 109.8, 106.8, 95.3, 56.1, 55.9, 54.5, 51.2, 46.1, 40.2, 32.1, 31.1, 29.8, 28.5, 28.5, 24.0, 21.9.

***N*-(2-Aminoethyl)-3-((3*S***,**6*S***,**12a*S*)-6-isobutyl-9-methoxy-1**,**4-dioxo-1**,**2**,**3**,**4**,**6**,**7**,**12**,**12a- octahydropyrazino[1’**,**2’:1**,**6]pyrido[3**,**4-*b*]indol-3-yl)propenamide (compound 4, figure 2a)**

To compound **3** (2.6 mg, 4.7 μmol) in dioxane (0.25 mL), 4 M HCl in dioxane (0.25 mL) was added. After stirring for 2 hours the solvent was removed to afford the desired product as an orange solid in a quantitative yield. LCMS *m/z* calc. for C_24_H_34_N_5_O_4_ [M+H]^+^ 456.3; found 456.3, *t*_R_ = 2.65. The product was used directly in the next step without further purification.

**6-(2-(4-((*E*)-2-(5**,**5-Difluoro-7-(thiophen-2-yl)-5*H*-4λ4**,**5λ4-dipyrrolo[1**,**2-*c*:2’**,**1’- *f*][1**,**3**,**2]diazaborinin-3-yl)vinyl)phenoxy)acetamido)-*N*-(2-(3-((3*S***,**6*S***,**12a*S*)-6-isobutyl-9- methoxy-1**,**4-dioxo-1**,**2**,**3**,**4**,**6**,**7**,**12**,**12a-octahydropyrazino[1’**,**2’:1**,**6]pyrido[3**,**4-*b*]indol-3- yl)propanamido)ethyl)hexanamide (compound 5, figure 2a)**

To compound **4** (1.15 mg, 2.3 μmol) in DCM (0.5 mL), DIPEA (2.0 μL, 11.6 μmol) and BODIPY-X- 630/650-SE (1.5 mg, 2.3 μmol) in DCM (0.5 mL) were added. The reaction mixture was stirred in the absence of light. A further 2.5 eq of DIPEA (1.0 μL, 5.8 μmol) was added after stirring for 2.5 hours and 30 μL of DMF after a further 1 hour. 5 eq of DIPEA (2.0 μL, 11.6 μmol) and DMF (100 μL) were added after an additional 2 hours and the mixture was stirred overnight. The solvent was removed, and the blue solid was purified by PTLC (5% MeOH in DCM). The product was further purified by semi-prep HPLC to afford the title product as a blue solid 0.2 mg, 9% LCMS *m/z* calc. for C_53_H_60_BF_2_N_8_O_7_S [M+H]^+^.= 1001.9; found 1001.7, *t*_R_ = 3.36 mins. HRMS (ESI-TOF) *m/z* calc. for C_53_H_60_BF_2_N_8_NaO_7_S [M+Na]^+^ 1023.4181; found 1023.4197 (Supplementary Figure 1). Analytical HPLC *t*_R_ = 19.83 mins, purity = 100%

**Supplementary figure 1:**
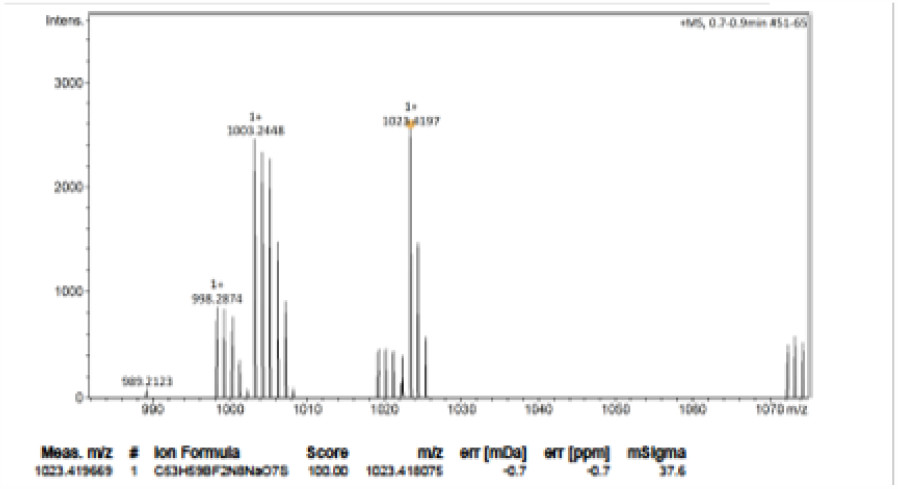
Mass spectrometry analysis of Ko143-X-BY630. High-resolution mass spectra (HRMS) were recorded on a Bruker microTOF mass spectrometer using MS electrcspray ionization operating in positive ion mode. Expected mass 1023.4181; found 1023.4197

**Supplementary figure 2:**
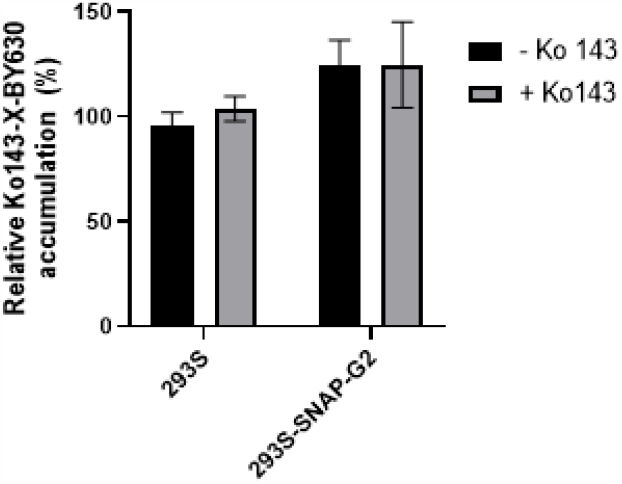
Ko143-X-BY630 is not transported by ABCG2. HEK293S cells or 293S-SNAP-ABCG2 were incubated with Ko143-X-BY630 in the presence or absence of Ko143. Cellular accumulation of Ko143-X-BY630 was determined by a fluorescence plate-reader and is expressed as a percentage of staining of HEK293S cells. Values represent the mean and standard error of three independent experiments.

## Notes

### Competing Interest Statement

The authors have declared no competing interest.

